# The cancer-promoting enzyme PKM2 binds RNA via a positively charged regulatory patch

**DOI:** 10.64898/2026.09.22.753476

**Authors:** Pia Sommerkamp, Christian Schillinger, Karine Lapouge, Antonio Biancolella, Dunja Ferring-Appel, Matthias W. Hentze

## Abstract

Pyruvate kinase M2 (PKM2) is a glycolytic enzyme that coordinates energy production with biosynthetic demands in physiologically proliferating and cancer cells. PKM2 has also emerged as a non-canonical RNA-binding protein. Here, we combine targeted PKM2 mutagenesis with RNA-protein interaction assays, biochemical and biophysical analyses to define how RNA binding relates to PKM2 allosteric control and oligomeric state. We show that the allosteric activator fructose-1,6-bisphosphate (FBP) strongly reduces PKM2 binding to RNA. Mutants with impaired oligomerization show reduced RNA binding, and RNA association is favored by the tetrameric state. A positively charged surface patch in the FBP-binding region is essential for RNA binding and displays emergent properties. Single-residue variants further link RNA association to FBP-responsive tetrameric conformations. Our data integrate RNA binding with PKM2 allosteric regulation.

## INTRODUCTION

RNA-binding proteins (RBPs) regulate virtually all steps of RNA metabolism, including RNA processing, localization, translation and decay. While classical RBPs typically contain dedicated globular RNA-binding domains (RBDs), proteome-wide RNA interactome studies have revealed a large number of non-canonical RBPs that lack such domains, and instead are primarily known for functions outside RNA biology, including metabolism, signaling, transport or protein homeostasis ^1–7^. This expansion of the RBPome has shifted attention from a unidirectional view, in which proteins regulate RNA fate, toward the concept of riboregulation, with RNA as a modulator of protein function. In this context, RNA binding may influence protein folding, enzymatic activity, ligand binding, protein-protein interactions, oligomerization, localization or higher-order assembly formation ^7^. This is particularly relevant for metabolic enzymes, many of which bind nucleotide-containing cofactors or metabolites, and may therefore harbor surfaces that can also engage RNA. The emerging challenge is to define the biological consequences of these RNA interactions, and how they intersect with the established biochemical regulation of the respective protein.

Pyruvate kinase M2 (PKM2) is a highly regulated glycolytic enzyme with central roles in proliferating cells and cancer metabolism ^8–11^. Pyruvate kinase catalyzes the final step of glycolysis, transferring a phosphate group from phosphoenolpyruvate (PEP) to ADP to generate pyruvate and ATP ^12^. The PKM gene encodes two isoforms, PKM1 and PKM2, through mutually exclusive alternative splicing of exon 9 or exon 10, respectively ^13,14^. Although PKM1 and PKM2 differ in only one exon, they display markedly different regulatory properties. PKM1 is constitutively active and is predominantly expressed in differentiated tissues, whereas PKM2 is allosterically regulated and enriched in embryonic, proliferating and cancer cells ^9,15–17^.

A defining feature of PKM2 regulation is its dynamic oligomeric equilibrium. Unlike PKM1, which is largely constitutively tetrameric and catalytically active, PKM2 interconverts between monomeric, dimeric, and tetrameric states, with the tetramer representing the highly active pyruvate kinase form ^10,18^. Tetramer formation is promoted by regulatory inputs, including FBP, a key physiological allosteric activator of PKM2 ^12,19^. FBP binds a regulatory pocket in the PKM2 C-domain, stabilizing the tetramer and shifting the equilibrium towards the active enzyme ^12,20–24^. Structural and biochemical studies have identified a regulatory patch including K433 and R436 as important for FBP binding and allosteric activation ^20^. This mechanism enables PKM2 to tune glycolytic flux to cellular metabolic demand: high PKM2 activity supports efficient conversion of PEP to pyruvate with ATP production, whereas reduced activity permits accumulation of upstream glycolytic intermediates that can be diverted into biosynthetic pathways, including nucleotide, amino acid, and lipid synthesis ^16,25^. In this way, PKM2 functions not only as a glycolytic enzyme but also as a metabolic control point that supports the metabolic flexibility of proliferating cells.

Beyond its canonical metabolic function, PKM2 has emerged as a non-canonical RBP. Several RNA interactome studies identified PKM2 among the metabolic enzymes that bind RNA in cells ^3,4,6,26^. Subsequent work suggested that PKM2 can interact with defined RNA targets and structures, linking PKM2 to post-transcriptional regulation and cancer-associated gene expression programs. For example, nuclear PKM2 was reported to bind and stabilize RNA G-quadruplex structures in precursor mRNAs, thereby promoting the expression of genes involved in cell migration and motility ^27^. In the cytoplasm, PKM2 associates with ER-associated ribosomes ^28^. Recently, cytoplasmic PKM2 was identified in association with ribosome-bound nascent chains through poly-ADP-ribosylated regions, and to promote translational pausing and decay of specific mRNAs in a metabolism-sensitive manner ^29^. These studies suggest that PKM2 may connect metabolic state with RNA regulation in multiple cellular compartments.

We recently identified over 100 significantly enriched PKM2-associated RNA regions in HeLa cells by soniCLIP ^30^. These included specific rRNA contacts that may mediate the reported ribosome association of PKM2. We also validated several mRNA targets by RIP-qRT-PCR, providing a defined set of PKM2-associated RNAs for functional and biochemical studies. However, the molecular basis of PKM2-RNA interactions and their consequences remain poorly understood. In particular, it is unclear how RNA binding intersects with the known regulatory features of PKM2 as an enzyme.

Here, we investigate the molecular determinants of PKM2-RNA interactions and their relationship to FBP responsiveness, oligomerization and enzymatic activity. We identify the positively charged, FBP-associated surface as a critical determinant of RNA binding, and show that RNA association is linked to the tetrameric state of PKM2. Single-residue analysis reveals that K433, R436 and R455 contribute to a regulatory patch with distinct contributions to RNA binding, FBP responsiveness and catalytic regulation. Thus, non-canonical RNA binding is embedded within the canonical allosteric architecture of PKM2.

## RESULTS

### A positively charged FBP-associated patch mediates PKM2 RNA binding

To gain molecular insight into the RNA-binding activity of PKM2, we first tested whether RNA binding is influenced by metabolites that interact with PKM2. Specifically, we assessed *in vitro* binding of PKM2 to two previously validated RNA targets ^30^ with increasing concentrations of its substrate PEP, its product pyruvate or the allosteric activator FBP by electrophoretic mobility shift assay (EMSA) (Figure 1A, Extended Data Fig 1A). EMSA analysis revealed that physiological concentrations of FBP, which are in the lower µM range (10-200 µM) ^31^, strongly reduced PKM2-RNA complex formation in a dose-dependent manner, whereas PEP caused only a mild reduction in RNA binding at high concentrations. In contrast, pyruvate did not detectably affect PKM2-RNA complex formation.

**Figure 1:**
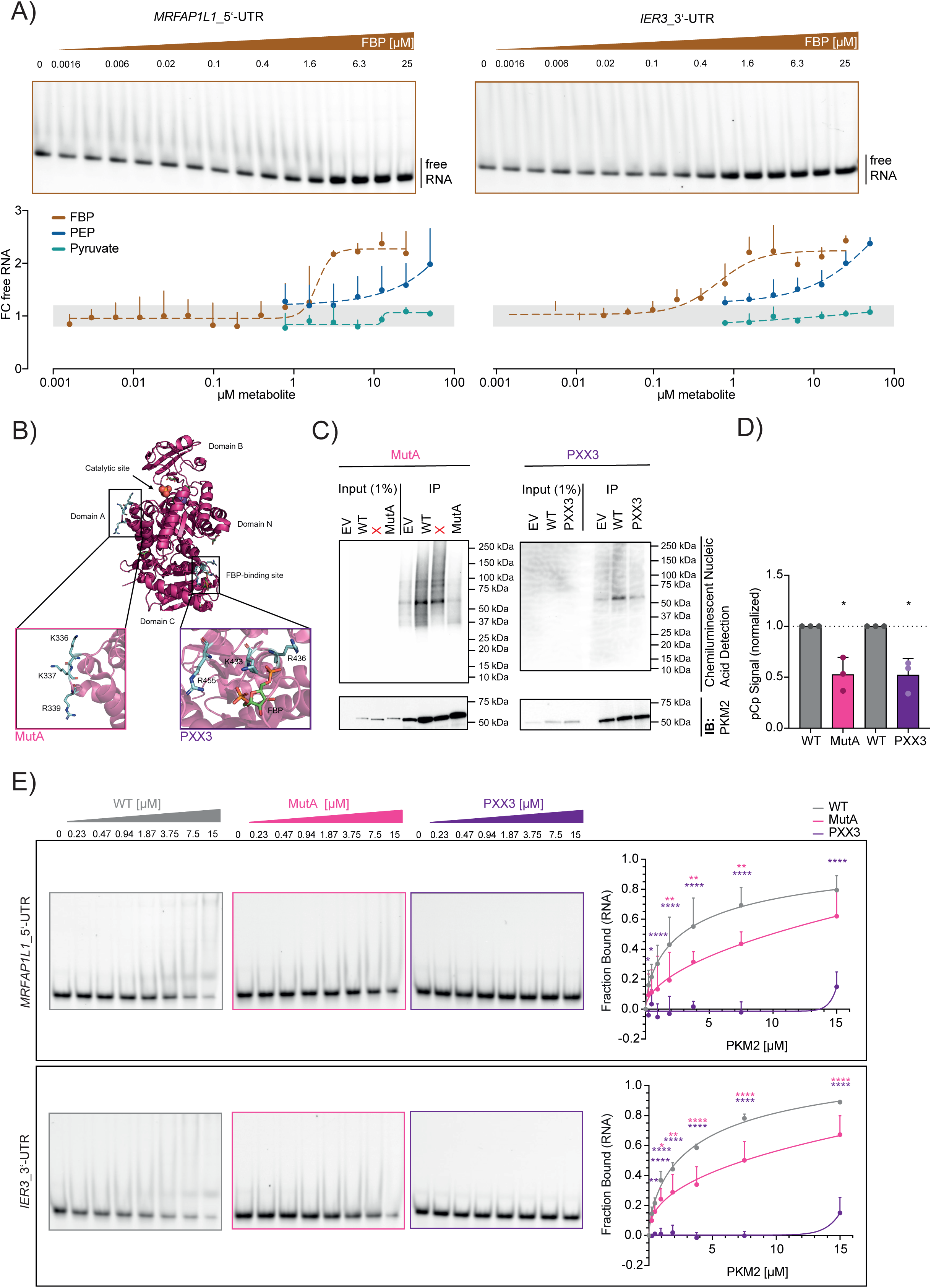
A positively charged FBP-associated patch mediates PKM2 RNA binding. (A) EMSA analysis and quantification of recombinant WT PKM2 and Cy5-labelled PKM2 RNA target oligos *MRFAP1L1*_5’-UTR (45mer) and *IER3*_3’-UTR (40mer) ^30^ in competition with its substrate phosphoenolpyruvate (PEP), its product pyruvate or the allosteric activator fructose-1,6-bisphosphate (FBP). N=3 (for each biological replicate an independent recombinant protein preparation was used). Mean +SD. Representative EMSA of FBP competition is shown. For PEP and pyruvate EMSAs see Extended Data Fig 1A. (B) Crystal structure of a single human wild-type PKM2 monomer within tetrameric association in complex with FBP. MutA: shows a loop of positively charged amino acid residues in domain A targeted to create the mutant MutA. PXX3: shows a positive patch within the FBP-binding region targeted to generate the mutant PXX3 and its respective single mutants. All panels: cartoon representation of PKM2 domains is shown in magenta, whereas mutated positions are highlighted as sticks in cyan. FBP is highlighted in stick representation with carbon, oxygen and phosphorus shown in green, red and orange, respectively. Images were created using PyMOL and the x-ray diffraction crystal structure 1T5A from the RCSB Protein Data Bank (https://doi.org/10.2210/pdb1t5a/pdb) ^12^. (C) pCp assay of WT, MutA and PXX3 overexpressed in HeLa cells. PKM2 was immunoprecipitated after crosslinking RNA-RBP complexes by UV-C light, cell lysis and RNA fragmentation. RNA was ligated to pCp-biotin and chemiluminescent nucleic acid detection was performed. Western blot analysis of PKM2 from the same experiment. X: analysis of an unrelated mutant. N=3. Representative blot is shown. (D) pCp quantification. Normalized to protein levels and WT overexpression control. N=3. Mean +SD. Paired student’s t test. *p < 0.05; **p < 0.01; ***p < 0.001; ****p < 0.0001. (E) EMSA analysis and quantification of recombinant WT PKM2, MutA and PXX3 and Cy5-labelled PKM2 RNA target oligos *MRFAP1L1*_5’-UTR (45mer) and *IER3*_3’-UTR (40mer) ^30^. N=3 (for each biological replicate an independent recombinant protein preparation was used). Mean +SD. Two-way ANOVA. *p < 0.05; **p < 0.01; ***p < 0.001; ****p < 0.0001. The color scheme indicates comparison between MutA or PXX3 with WT PKM2, respectively. Representative EMSAs are shown.

FBP binds to a regulatory pocket in PKM2 involving K433 and neighboring residues, thereby stabilizing the active tetrameric state and enhancing pyruvate kinase activity ^12,20,21,23,24^. We noted that this region forms part of a positively charged surface patch (Extended Data Fig 1B-C), suggesting that the antagonism of FBP towards RNA binding may involve a region overlapping with, or in close proximity to, the FBP-binding site. To test this possibility, we generated a PKM2 mutant in which three positively charged residues within this patch were substituted by alanine (PXX3: K433A, R436A, R455A) (Figure 1B). In parallel, to distinguish effects on RNA binding from potential contributions of PKM2 oligomerization, we generated a second mutant targeting basic residues in the dimerization domain A (MutA: K336A, K337A, R339A) (Figure 1B).

Both PKM2 mutants were overexpressed in HeLa cells such that the endogenous PKM2 background signal (empty vector, EV) is low in comparison to the exogenous expression of wild type (WT) protein and mutants, respectively; their RNA-binding activity was analyzed using the pCp assay (Figure 1C-D)^32^. Compared with exogenous WT PKM2, both MutA and PXX3 showed significantly reduced RNA binding *in cellulo*, suggesting that both oligomerization and the positively charged FBP-associated patch contribute to the PKM2-RNA interaction.

To follow-up on these findings, we assessed the RNA-binding activity of recombinant MutA and PXX3 proteins *in vitro* by EMSA. Consistent with the cellular pCp assays, both mutants displayed impaired RNA binding (Figure 1E). However, the extent of the impairment differed markedly between the two mutants: while MutA showed a moderate reduction in binding to specific PKM2 RNA targets, RNA binding by PXX3 was almost completely abolished. Together, these results indicate that PKM2-RNA interactions depend critically on a positively charged surface surrounding the FBP-binding region, whereas disruption of the dimerization interface in MutA impairs RNA binding to a lesser extent.

### PKM2 RNA binding requires an intact FBP-responsive surface and is favored by the tetrameric state

Next, we performed a detailed biophysical and enzymatic characterization of WT PKM2, MutA and PXX3, with particular focus on their stability, oligomerization state and responsiveness to FBP. We first used nano differential scanning fluorimetry (nanoDSF) to assess the thermal stability of recombinant WT PKM2, MutA and PXX3 under ‘stabilizing’ protein buffer conditions and in a ‘destabilizing’ PBS-based buffer (Figure 2A, Extended Data Fig 2A-B). In protein buffer, WT PKM2 and PXX3 displayed melting temperatures of approximately 62 °C, whereas MutA showed a minor reduction in thermal stability. Under destabilizing PBS conditions, the melting temperatures of WT PKM2 and PXX3 were reduced to approximately 60 °C, while MutA showed no further decrease. These results suggest that mutations in the dimerization domain impair the stabilizing interactions required for optimal PKM2 oligomerization, which is reflected by reduced thermal stability under otherwise stabilizing buffer conditions. Addition of FBP did not markedly alter the quantified melting temperatures (Figure 2A, Extended Data Fig 2A). However, analysis of the nanoDSF thermograms revealed a second transition at approximately 70 °C upon FBP treatment of WT PKM2 and MutA, suggesting stabilization of a minor protein subfraction that was not captured by the main melting-temperature quantification (Figure 2A, Extended Data Fig 2A). In contrast, no comparable FBP-induced shift was observed for PXX3. These data indicate that PXX3, but not MutA, has impaired responsiveness to FBP, consistent with previous reports implicating the region surrounding K433 in FBP binding ^20^.

**Figure 2:**
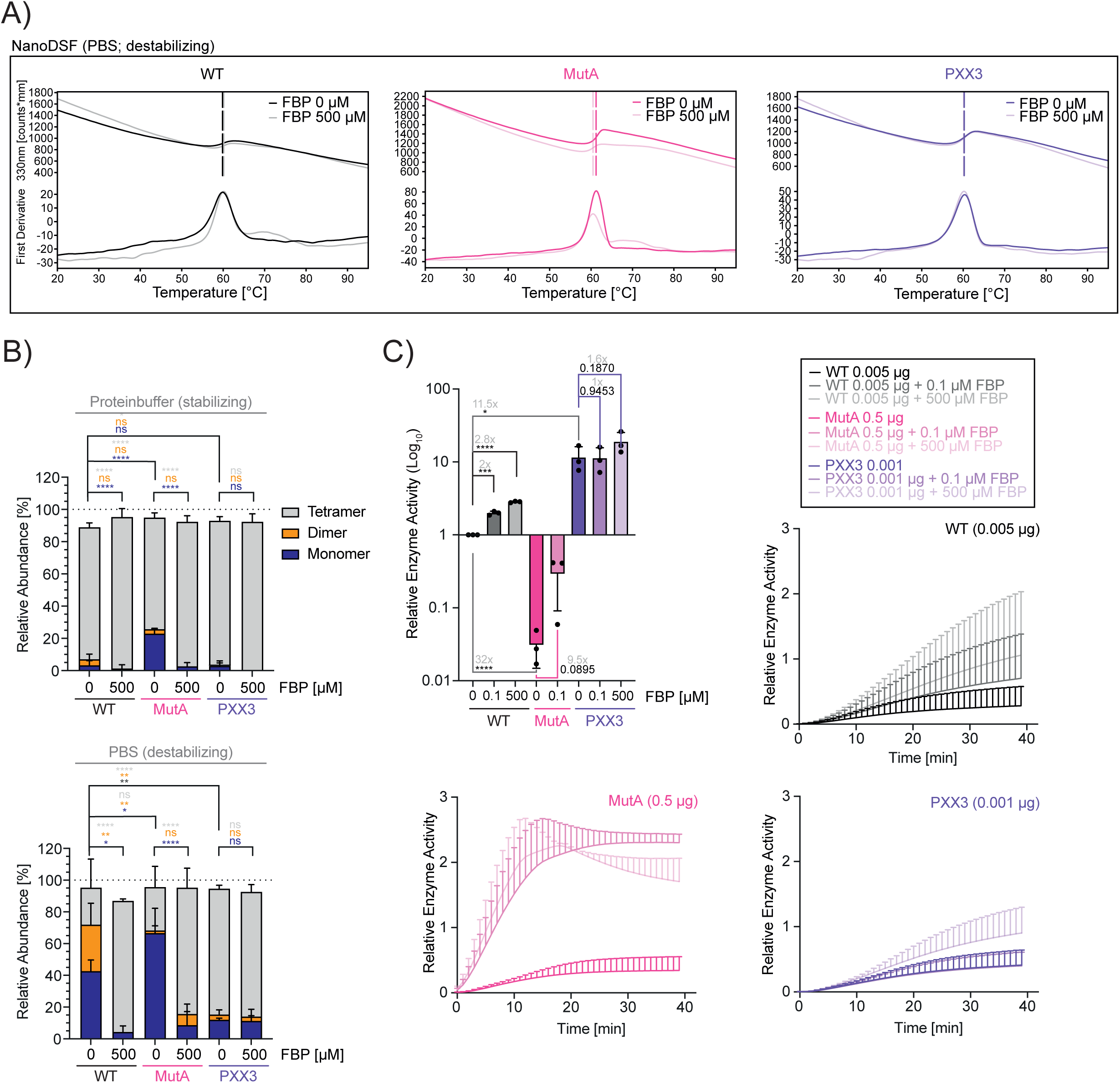
PKM2 RNA binding requires an intact FBP-responsive surface and is favored by the tetrameric state. (A) Representative NanoDSF thermograms of 3 µM PKM2 WT, MutA and PXX3 ±500 µM FBP in PBS + 2.5 mM MgCl_2_. N=3 (for each biological replicate an independent recombinant protein preparation was used). (B) Mass photometry quantification of normalized counts of PKM2 WT, MutA and PXX3 in protein buffer or PBS +2.5 mM MgCl_2_ ±FBP. N=3 (for each biological replicate an independent recombinant protein preparation was used). Mean +SD. Two-way ANOVA. *p < 0.05; **p < 0.01; ***p < 0.001; ****p < 0.0001; ns: not significant. The color scheme indicates comparison between different oligomerization states. (C) Quantification of pyruvate kinase activity levels normalized to WT and pyruvate kinase activity curves ±FBP. In order to stay within the assay kit’s linearity capacities, 0.005 µg WT PKM2, 0.5 µg MutA and 0.001 µg PXX3 were used. Different amounts of recombinant protein were taken into account and corrected during calculation of relative enzyme activity. MutA + 500 µM FBP was not quantified, as it was outside of the assay’s linear range. N=3 (for each biological replicate an independent recombinant protein preparation was used). Mean +SD. Unpaired student’s t test. *p < 0.05; **p < 0.01; ***p < 0.001; ****p < 0.0001. Fold changes of relative enzyme activity are indicated in gray.

To directly determine how these mutations affect PKM2 oligomerization, we next performed mass photometry under both stabilizing and destabilizing buffer conditions (Figure 2B, Extended Data Fig 2C). Under stabilizing protein buffer conditions, WT PKM2 and PXX3 were predominantly tetrameric. In contrast, MutA showed impaired tetramer formation, with a significantly increased monomer fraction, as expected. FBP had only minor effects on WT PKM2 and PXX3 under these conditions, likely because both proteins were already largely tetrameric. By contrast, FBP shifted MutA toward the tetrameric state, indicating that MutA retains the ability to respond to FBP and that FBP can stabilize tetramers formed by this mutant.

Under destabilizing PBS conditions, WT PKM2 showed a pronounced reduction in the tetrameric fraction, accompanied by an increase in monomeric and dimeric species. In line with its known role as an allosteric activator, FBP efficiently restored tetramer formation of WT PKM2. MutA was highly sensitive to destabilizing conditions and was predominantly monomeric in PBS. Nevertheless, FBP again promoted tetramer formation, further supporting the conclusion that MutA is defective in oligomerization but not in FBP responsiveness. PXX3 remained predominantly tetrameric even under destabilizing conditions, and FBP did not further alter its oligomeric distribution. Thus, the PXX3 mutation of the positively charged patch renders PKM2 insensitive to FBP and strongly impairs RNA binding, while stabilizing the tetrameric state.

Because PKM2 tetramer formation is closely linked to enzymatic activity, we next measured the pyruvate kinase activity of WT PKM2, MutA and PXX3 in the absence and presence of a low or high concentration of FBP (Figure 2C). Recombinant WT PKM2 displayed the expected enzymatic activity, which was further increased in response to FBP. MutA showed strongly reduced enzymatic activity, consistent with its impaired tetramerization. However, FBP treatment partially rescued MutA activity, in agreement with the mass photometry data showing FBP-dependent stabilization of MutA tetramers. At high FBP concentrations, MutA activity was further increased, although precise quantification was limited because the assay exceeded the linear range. In contrast, PXX3 displayed markedly elevated basal enzymatic activity, approximately 11.5-fold higher than WT PKM2 under steady-state conditions. This hyperactivity is consistent with the highly stable tetrameric state observed for PXX3. FBP did not further increase PXX3 activity, supporting the conclusion that this mutant is functionally unresponsive to FBP.

Taken together, these data reveal that MutA and PXX3 impair PKM2 RNA binding by distinct mechanisms. MutA displays defective oligomerization, increased monomer formation, and reduced enzymatic activity, all of which can be partially rescued by FBP. Since FBP binds and stabilizes the PKM2 tetramer ^12,20,21,23,24^, and since RNA and FBP functionally compete for binding to PKM2, the reduced RNA-binding activity of MutA likely reflects reduced availability of an RNA binding-competent tetrameric conformation. This suggests that monomeric or dimeric PKM2 are less competent for RNA binding, similar to what has been reported for FBP binding ^22^. PXX3, in contrast, forms a highly stable and hyperactive tetramer, lacks detectable responsiveness to FBP, and shows a near-complete loss of RNA binding. Thus, the positively charged patch mutated in PXX3 is not required for tetramer formation itself, but is essential for the interaction with both FBP and RNA.

We also tested whether RNA targets reciprocally affect PKM2 oligomerization or enzymatic activity. Under the conditions examined, we did not obtain reproducible effects of RNA on PKM2 activity or oligomerization. Thus, while our data support a model in which RNA binding is coupled to the FBP-responsive tetrameric state, whether RNA directly modulates the PKM2 oligomerization state and/or enzymatic activity remains unresolved.

### Single residue mutations uncouple PKM2 RNA binding, FBP responsiveness and enzymatic activity

To determine whether RNA and FBP engage the same residues within the positively charged patch, or whether they depend on overlapping but distinct determinants, we next analyzed the contribution of each mutated residue of PXX3 individually. We therefore generated single alanine-substitution mutants of K433, R436, and R455, all of which are located within the FBP-associated positively charged patch disrupted in PXX3.

We first assessed RNA binding of the single mutants by EMSA (Figure 3A). Mutation of all three residues individually reduced RNA-binding, but to different extents. R436A and R455A showed significantly impaired, but still detectable, RNA binding. In contrast, K433A almost completely lost RNA binding and thereby most closely phenocopied the PXX3 triple mutant in this regard. These data indicate that K433 is critical for the PKM2-RNA interaction, while R436 and R455 contribute to RNA binding.

**Figure 3:**
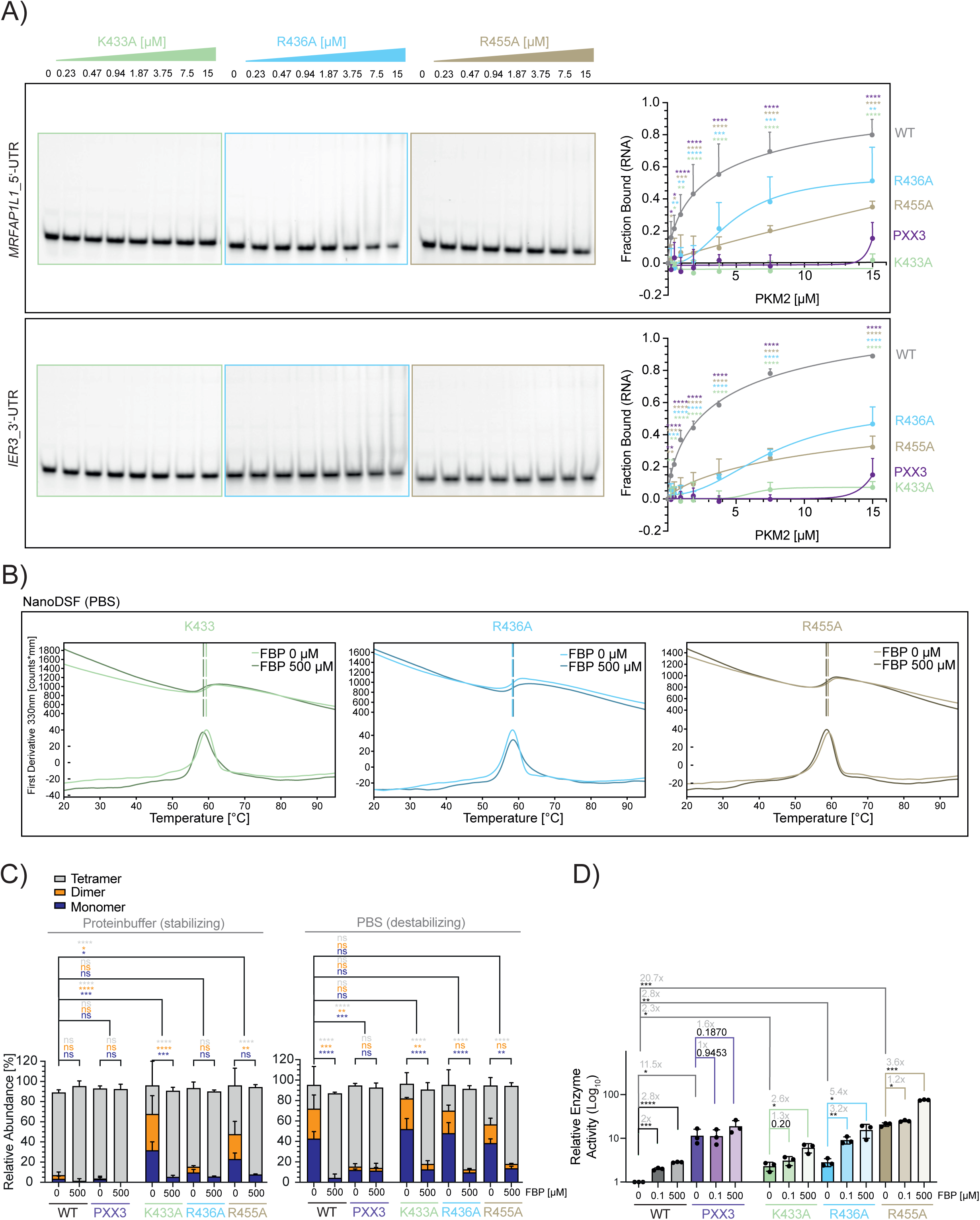
Single residue mutations uncouple PKM2 RNA binding, FBP responsiveness and enzymatic activity. (A) EMSA analysis and quantification of recombinant K433A, R436A and R455 and Cy5-labelled PKM2 RNA target oligos *MRFAP1L1*_5’-UTR (45mer) and *IER3*_3’-UTR (40mer) ^30^. WT and PXX3 are shown for comparison (see Figure 1E). N=3 (for each biological replicate an independent recombinant protein preparation was used). Mean +SD. Two-way ANOVA. *p < 0.05; **p < 0.01; ***p < 0.001; ****p < 0.0001. The color scheme indicates comparison between single mutants with WT PKM2, respectively. Representative EMSAs are shown. (B) Representative NanoDSF thermograms of 3 µM K433A, R436A and R455A ±500 µM FBP in PBS +2.5 mM MgCl_2_. N=3 (for each biological replicate an independent recombinant protein preparation was used). (C) Mass photometry quantification of normalized counts of K433A, R436A and R455A in protein buffer or PBS +2.5 mM MgCl_2_ ±FBP. WT and PXX3 are shown for comparison (see Figure 2B). N=3 (for each biological replicate an independent recombinant protein preparation was used). Mean +SD. Two-way ANOVA. *p < 0.05; **p < 0.01; ***p < 0.001; ****p < 0.0001; ns: not significant. The color scheme indicates comparison between different oligomerization states. (D) Quantification of pyruvate kinase activity levels normalized to WT FBP. In order to stay within the assay kit’s linearity capacities, 0.001 µg K433A, R436A and R455A were used, respectively. Different amounts of recombinant protein were taken into account and corrected during calculation of relative enzyme activity. WT and PXX3 are shown for comparison (see Figure 2C). N=3 (for each biological replicate an independent recombinant protein preparation was used). Mean +SD. Unpaired student’s t test. *p < 0.05; **p < 0.01; ***p < 0.001; ****p < 0.0001. Fold changes of relative enzyme activity are indicated in gray.

To understand if the strong RNA-binding defect of K433A reflects direct loss of a critical RNA-contact residue or instead results from altered FBP responsiveness and oligomerization, we analyzed thermal stability and FBP responsiveness by nanoDSF (Figure 3B, Extended Data Fig 3A). In contrast to WT PKM2 (Figure 2A), all single mutants showed reduced responsiveness to FBP, as reflected by the loss or marked reduction of the FBP-induced secondary transition at ∼70 °C observed in the nanoDSF thermograms. This effect was most pronounced for R455A, for which no clear FBP-induced secondary transition was detected under either stabilizing or destabilizing buffer conditions. R436A, and potentially K433A, however, retained a weak secondary transition around 65 °C, indicating residual interaction with FBP. These findings are consistent with previous reports identifying K433 as an important residue for FBP binding ^20^, whereas R436 appears to make a smaller contribution.

We then directly assessed the oligomeric state of the single mutants by mass photometry (Figure 3C, Extended Data Fig 3B). Interestingly, none of the single mutants reproduced the oligomerization phenotype of PXX3. Whereas PXX3 formed a highly stable tetramer unresponsive to FBP, K433A and R455A showed a pronounced shift toward monomeric and dimeric species under stabilizing protein buffer conditions. Under PBS buffer conditions, K433A and R455A showed a further reduction in the tetrameric fraction, although their overall distribution was now comparable to that of WT PKM2. In contrast, R436A behaved more similarly to WT PKM2 even in the stabilizing protein buffer, showing only a modest increase in the monomer fraction while predominantly forming tetramers. Importantly, all three single mutants remained responsive to high-dose FBP in mass photometry. Of note, FBP responsiveness seems to be assay-dependent, with nanoDSF and mass photometry capturing distinct aspects of FBP-induced stabilization and oligomeric redistribution. FBP treatment shifted the oligomeric equilibrium toward the tetrameric state and strongly reduced the monomeric and dimeric fractions, both under stabilizing and destabilizing buffer conditions. Thus, although K433A and R455A show reduced FBP responsiveness in nanoDSF, their oligomerization defects can still be rescued by high-dose FBP. This distinguishes the single mutants from PXX3, which forms an FBP-insensitive, highly stable tetramer.

Next, we examined how these changes affect PKM2 catalytic activity (Figure 3D, Extended Data Fig 3C). K433A and R436A showed mildly increased basal enzymatic activity compared with WT PKM2, despite the altered oligomeric distribution observed for K433A. R436A remained responsive to low-dose FBP, resulting in a further increase in enzymatic activity, consistent with its residual FBP responsiveness in nanoDSF and near-WT oligomerization behavior in mass photometry. In contrast, K433A responded poorly to physiological FBP concentrations, supporting the central role of K433 in FBP-mediated activation. R455A displayed a markedly different activity pattern: despite its impaired oligomerization profile in mass photometry compared to PXX3, R455A showed strongly elevated basal enzymatic activity (20.7x), similar to the hyperactive PXX3 mutant. This discrepancy may reflect differences between the conditions used for mass photometry and activity assays, especially with regard to buffer conditions, or a catalytically competent conformation not fully captured by static oligomer distributions. Moreover, like K433A and PXX3, R455A showed little response to physiological FBP levels. These findings suggest that mutation of R455 is a major contributor to the hyperactive enzymatic phenotype observed in PXX3 and reveals a partial uncoupling between catalytic activity and the oligomeric distribution detected under these assay conditions.

Together, the single-mutant analyses reveal that the multi-modal changes of the PXX3 mutant cannot be attributed to any of the three mutated residues alone (Table 1). K433A most closely mimics the RNA-binding defect of PXX3, but unlike PXX3 it does not form a hyperstable tetramer. Instead, K433A exhibits a strong shift toward monomeric and dimeric species, suggesting that its loss of RNA binding may at least partly result from reduced availability of the RNA-competent tetrameric state. This interpretation is supported by the milder phenotype of R436A, which retains substantial tetramer formation, residual FBP responsiveness, and only partially impaired RNA binding. By contrast, R455A contributes strongly to FBP-dependent regulation and enzymatic activity, as it attains a hyperactive enzymatic state similar to PXX3, despite an apparent destabilization of tetramer formation. Thus, K433, R436, and R455 contribute differently to RNA binding, FBP responsiveness, oligomerization and enzymatic activity, and all three amino acids exert effects on the above parameters. These data support a model in which productive PKM2-RNA interaction requires both an intact positively charged surface and the appropriate tetrameric conformation, while the combined disruption of this patch in PXX3 generates an FBP-insensitive, RNA-binding-deficient, hyperactive tetramer.

**Table 1:** Overview of PKM2 RNA-binding mutants and associated phenotypes.

|  | WT | MutA | PXX3 | K433A | R436A | R455A |
| --- | --- | --- | --- | --- | --- | --- |
| Mutation(s) |  | K336A<br>K337A<br>R339A | K433A<br>R436A<br>R455A | K433A | R436A | R455A |
| Position of Mutation |  | (+)Loop<br>dimerization<br>domain A | (+)Patch<br>encompassing<br>FBP-binding<br>site | Reported to be<br>main AA<br>involved in FBP<br>binding | Reported to be<br>involved in FBP<br>binding |  |
| RNA-binding |  | ↓ | ↓↓↓ | ↓↓↓ | ↓ | ↓↓ |
| Tetramer formation |  | ↓ | ↑↑ | ↓↓ | unchanged | ↓ |
| Enzymatic activity |  | ↓↓↓ | ↑↑↑ | ↑ | ↑ | ↑↑↑ |
| FBP-sensitivity EA<br>(100 nM) | ++ | +++ | - | - | ++ | - |
| FBP-sensitivity<br>NanoDSF (500 μM) | ++ | ++ | - | + | + | - |
| FBP-sensitivity MP<br>(500 μM) | ++ | ++ | - | ++ | ++ | ++ |
| Oligomerization |  | Oligomerization<br>impaired | Highly stable<br>tetramer | Oligomerization<br>impaired | Oligomerization<br>impaired | Oligomerization<br>impaired |
| FBP-responsiveness |  | High,<br>oligomerization<br>defect can be<br>rescued by FBP | Absent | Impaired FBP-responsiveness | mostly<br>unchanged | Impaired FBP-responsiveness |
| RNA-binding mode<br>\| Hypothesis | Binding<br>depends on<br>presence of<br>tetramer<br>→ RNA binds to<br>(+)Patch in<br>absence of FBP | Less tetramer<br>→ Indirect<br>RNA-binding<br>mutant | No (+)Patch<br>→ Complete<br>loss of RNA-binding activity | Less tetramer<br>→ Indirect<br>RNA-binding<br>mutant<br>OR<br>Loss of<br>essential (+)<br>charge<br>→ Complete<br>loss of RNA-binding activity | Loss of (+)<br>charge<br>→ Slightly<br>reduced RNA-binding activity | Less tetramer<br>→ Indirect<br>RNA-binding<br>mutant |

## DISCUSSION

In this study, we define molecular determinants of RNA binding by PKM2, a glycolytic enzyme previously identified as a non-canonical RBP ^3,4,6,26^. The molecular basis of PKM2-RNA binding has remained unclear, and our data address this gap by linking PKM2-RNA interaction to two central features of PKM2 regulation: FBP-dependent allosteric control and oligomeric state. This places PKM2 among a growing class of non-canonical RBPs for which RNA binding must be considered in the context of protein conformation, stability and assembly state ^33–36^, and cannot be understood solely from primary sequence or isolated surface charge.

The allosteric regulator FBP strongly reduces binding of PKM2 to RNA targets at physiological concentrations, suggesting that PKM2-RNA interactions and their functional consequences in cells may be regulated metabolically by FBP. FBP binds to an allosteric pocket in the C-domain and stabilizes the active tetrameric enzyme, thereby increasing pyruvate kinase activity ^12,20–24^. Residues in this region, including K433 and R436, have been implicated in FBP binding and allosteric activation ^20,22^. Our findings therefore suggest that PKM2-associated RNAs engage a surface that overlaps with, lies adjacent to, or is conformationally coupled to the FBP-binding pocket. This complements observations for other metabolic RBPs, where RNA binding can involve substrate- or cofactor-binding regions, as described for GAPDH, ENO1 and other nucleotide-binding metabolic enzymes such as SHMT1 ^33–36^. Thus, RNA binding by PKM2 could be embedded in a pre-existing allosteric regulatory network rather than represent an independent accessory function.

The FBP-associated surface contains a positively charged patch including K433, R436 and R455. Positively charged protein surfaces can engage the negatively charged RNA backbone and are frequently implicated in RNA binding by non-canonical RBPs lacking classical RNA-binding domains ^37–39^. Nonetheless, our data argue that this region is not merely an electrostatic interaction surface for RNA. PKM2 binds specific RNA targets identified in cells, whereas GC content- and size-matched control RNAs show little or no interaction in EMSA experiments ^30^. Mutation of the positive patch in PXX3 strongly impaired RNA binding in cells and *in vitro*, abolished detectable FBP responsiveness and, unexpectedly, generated a hyperstable, hyperactive tetramer. Thus, this region appears to function as a regulatory interface of PKM2 at which RNA binding, metabolite sensing, oligomerization and catalytic activity converge.

MutA and PXX3 define two separate routes to impaired RNA binding. MutA, which carries mutations in the dimerization domain, showed increased monomer formation, reduced tetramer abundance and strongly decreased enzymatic activity. These defects were partially rescued by FBP, indicating that MutA remains FBP-responsive. Thus, it can be concluded that impaired oligomerization correlates with reduced RNA binding. According to our proposed model, MutA reduces RNA binding indirectly by limiting the availability of the RNA-competent PKM2 tetramer.

PXX3 showed a distinct molecular phenotype. Despite forming a highly stable tetramer, PXX3 exhibited a near-complete loss of RNA binding and lacked detectable FBP responsiveness. This implies that tetramerization alone is not sufficient for RNA binding; rather, an intact positive patch surrounding the FBP-binding site is also required. Unexpectedly, PXX3 formed a hyperstable and hyperactive tetramer, partially reminiscent of the constitutively active behavior of PKM1. However, PXX3 is not simply a PKM1-like variant: it combines three alanine substitutions within the FBP-associated positive patch and lacks detectable FBP responsiveness or RNA binding. In this respect, PXX3 differs from previously described PKM1-mimetic PKM2 mutants such as C424L, in which substitution of the PKM2-specific cysteine with the PKM1-like leucine promotes tetramer formation in the absence of an allosteric activator ^40^. Whereas C424L highlights how individual isoform-specific residues can shift PKM2 toward a PKM1-like tetrameric state, PXX3 reveals a different regulatory outcome: a hyperactive tetramer that has lost both FBP responsiveness and RNA-binding competence. Since the catalytic center of PKM2 is distant from the mutated region, the increased activity of PXX3 is most consistent with an altered quaternary structure or allosteric state rather than a direct effect on catalytic chemistry. This interpretation is in line with structural work showing that PKM2 activity depends not only on tetramer abundance, but also on transitions between inactive T-state and active R-state tetrameric conformations ^41,42^. FBP stabilizes the active R-state, whereas the patient-derived K422R mutant, for example, forms a stable but catalytically impaired T-state tetramer ^43^. PXX3 may therefore bias PKM2 toward an active, R-like conformation that partially mimics the FBP-bound state, although structural analyses will be required to test this directly. Taking into account that FBP inhibits RNA binding, the loss of RNA binding by PXX3 may also be explained by structural mimicry of the FBP-bound state.

Dissecting PXX3 further, the single-residue mutants indicate that this surface acts as a coupled regulatory region with emergent properties rather than as a set of residues with additive effects. Published PKM2 mutant and post-translational modification studies showed that local perturbations near the FBP-binding region can have non-local effects on FBP responsiveness, oligomerization and catalytic output ^20,44^. K433A most closely reproduced the RNA-binding defect of PXX3, consistent with the established role of K433 as a central FBP-contact residue ^20^. R436A showed the mildest phenotype, suggesting that it contributes to the electropositive surface without acting as a dominant allosteric node. R455A, in contrast, retained detectable RNA binding and showed strongly elevated basal enzymatic activity, indicating that R455 contributes prominently to allosteric regulation. The fact that none of the single mutants fully reproduced PXX3 indicates that the triple mutant is not simply the additive consequence of losing three positive charges. Instead, PXX3 appears to display emergent properties, in which simultaneous mutation of K433, R436 and R455 creates a regulatory state that differs qualitatively from each individual substitution. Together, these data support an integrated surface model in which K433, R436 and R455 collectively tune RNA binding, FBP responsiveness, oligomerization and catalytic output.

An important open question is how PKM2 recognizes its RNA targets. The absence of a simple enriched RNA sequence motif from the identified RNA targets ^30^ suggests that PKM2 may recognize RNA structure, local RNA context, or broader physicochemical features rather than a short linear sequence. This may help reconcile the different RNA-related activities reported for PKM2, including nuclear binding to RNA G-quadruplex structures and cytoplasmic ribosome-associated interactions ^27–29^. While we also identified rRNA-derived regions as specific PKM2 targets ^30^, we selected mRNA-derived targets for the experiments using synthetic RNAs reported here to avoid ambiguities in interpretation based on the extensive modification of rRNA. Nuclear and cytoplasmic PKM2 differ in associated functions, binding partners, metabolite environments and oligomeric states ^18^. PKM2-RNA binding may therefore represent a set of context-dependent interactions shaped by RNA structure, subcellular localization and PKM2 conformation allowing for compartment-specific regulation.

Whether PKM2-RNA interactions primarily reflect a moonlighting function toward bound RNAs or riboregulation of PKM2 activity, or both, remains to be resolved. A moonlighting role would imply that PKM2 affects target-RNA fate. A riboregulatory model would entail that RNA modulates PKM2 function, for example by affecting FBP binding, tetramer stability or enzymatic activity. Because RNA and FBP functionally compete *in vitro*, RNA-mediated regulation of PKM2 remains an attractive possibility. However, we explored several *in vitro* assays that did not yield conclusive effects of RNA on PKM2 enzymatic activity. Such effects may require specific RNA structures, metabolite concentrations, post-translational modifications or cellular states that are not fully captured by the experimental conditions that we tested. The interpretation of experiments expressing our PKM2 mutants in transfected cells was confounded by the formation of mixed tetramers with endogenous PKM2. Thus, follow-up studies involving mutagenesis of endogenous PKM2 should yield answers to these questions.

Post-translational modifications may provide an additional regulatory layer ^18^. K433 acetylation has been implicated in promoting PKM2 nuclear localization of the enzymatically inactive dimeric PKM2 ^21,45^. R455 has also been linked to regulatory control of PKM2 and has been reported as a site of arginine methylation ^46^. Since our data identify these amino acids as important for RNA binding, modifications at or near the FBP-associated positive patch may themselves control regulatory effects by RNA. This possibility is particularly relevant in proliferating and cancer cells, where PKM2 regulation is extensively remodeled ^8–11^.

Limitations of this study should also be considered. EMSA provides comparative RNA-binding information, but the coupled equilibria of RNA binding, oligomerization and ligand responsiveness complicate quantitative interpretations. Recombinant protein assays may not fully reproduce cellular contexts, where metabolites, post-translational modifications, binding partners and compartment-specific conditions influence PKM2. Conversely, cellular assays may be affected by endogenous PKM2 and mixed oligomer formation with overexpressed variants. Future approaches combining defined oligomeric states, quantitative binding measurements and structural analysis will be needed to distinguish direct RNA contacts from indirect conformational effects.

In conclusion, our data identify the FBP-associated positive patch as a regulatory interface linking PKM2 RNA binding to tetramer formation and allosteric control. Rather than being mediated by a single residue, PKM2-RNA interactions depend on the combined contribution of this surface and an RNA-competent tetrameric state. This places RNA binding within the established regulatory architecture of PKM2 and suggests that RNA, metabolites, oligomerization and enzymatic activity are interconnected features of PKM2 function. In this context, PKM2 also serves as a model for unraveling how non-canonical RNA binding can be integrated into the regulation of metabolic enzymes. Future work will determine whether this interaction primarily affects target-RNA fate or PKM2 function, or both.

## MATERIALS AND METHODS

### Cell Culture

HeLa cells were cultured in high-glucose (4.5 g/L D-glucose) Dulbecco’s modified Eagle’s medium (DMEM) supplemented with 2 mM L-glutamine, 10% heat-inactivated fetal bovine serum (FBS) (10270106, Thermo Fisher Scientific) and 100 U/mL penicillin-streptomycin in an incubator at 37 °C and 5% CO_2_.

### Overexpression of PKM2 Mutants

250,000 HeLa cells were seeded per well in 6-well plates containing 1.7 mL of high-glucose DMEM medium supplemented with 2 mM L-glutamine, 10% FBS (10270106, Thermo Fisher Scientific) and 100 U/mL penicillin-streptomycin per well and grown for 3 h at 37 °C and 5% CO2. Transfection was performed using the Lipofectamine^TM^ 3000 transfection kit (Invitrogen), following the manufacturer’s instructions with adaptations. 1 μg of plasmid DNA (pcDNA3.1+/C-(K)-DYK expression vectors, see supplemental material vector maps for details), 1 μL of Lipofectamine^TM^ 3000 and 8 μL of P3000^TM^ were used per well and condition. Treated cells were incubated at 37 °C and 5% CO2. Medium was exchanged 24 h post transfection. Crosslinking and cell lysis were performed after 48 h (*see pCp-Assay*)

### pCp-Assay

#### Generation of cross-linked HeLa cell lysates

HeLa cells were cultured in 15 cm culture dishes until 80-90% confluent. Plates were placed on ice and washed twice with ice-cold PBS. Remaining PBS was removed completely and plates were left to dry upside down for 30 s. While on ice, cells were cross-linked using the SpectroLinker™ XL-1500 UV-Crosslinker (150 mJ/cm^2^ UV radiation). At least one plate per condition was kept as the non-cross-linked control. Immediately after cross-linking, cells were lysed and scraped in 500 µL of ice-cold iCLIP100 lysis buffer (see Buffers, supplemental material) per plate. Lysates were vortexed every 5 min for 15 min and kept on ice in between. Lysates were sonicated using a Diagenode sonicator Bioruptor® Plus (5 cycles; 30 seconds ON, 30 seconds OFF; high power mode at 4 °C). Samples were centrifuged at 13,400 rcf for 10 min at 4 °C. Supernatants were transferred to fresh tubes. Protein concentration was measured using the Qubit™ Protein Broad Range (BR) Assay Kit on a Qubit™ 4 Fluorometer.

#### pCp procedure

Protein A magnetic beads (Dynabeads™, Bio-Rad) were used for immunoprecipitation of rabbit antibodies. For each IP reaction, 20 µL of beads were used. Beads were washed three times with 1 mL of iCLIP lysis buffer (see Buffers, supplemental material), followed by resuspension in 100 µL lysis buffer per IP. Antibodies added: PKM2 (D78A4 Cell Signaling, 1 µg/IP), rabbit IgG (Invitrogen #10500C, 1 µg/IP). Beads and antibodies were incubated on a rotating wheel at 13 rpm in a cold room (4 °C) for 2 h or overnight. Cell lysates (80 - 500 µg total protein per sample) were adjusted to 500 µL using iCLIP lysis buffer. DNase digestion was performed using 2 µL Turbo DNase (Thermo Fisher) at 37 °C for 5 min at 850 rpm. RNA was fragmented by sonication using a Diagenode Bioruptor™ Plus sonicator (20 cycles; 30 seconds ON, 30 seconds OFF; high power mode at 4 °C). A 5 µL aliquot was kept as input control. Following bead-antibody coupling, beads were washed three times with 1 mL iCLIP lysis buffer and resuspended in 50 µL iCLIP lysis buffer per IP. Beads were added to each digested lysate and incubated at 4 °C for 2 h on a rotating wheel.

Each sample was washed twice with 900 µL of ice-cold iCLIP100 lysis buffer, twice with 900 µL of ice-cold high salt wash buffer (see Buffers, supplemental material) without and once with LiCl (see Buffers, supplemental material) and once with 500 µL of ice-cold wash buffer (see Buffers, supplemental material) (rotating, 3 min per wash step). While separated on the magnetic rack, the samples were resuspended in 500 µL of ice-cold wash buffer, to which 500 µL of FastAP buffer (see Buffers, supplemental material) were added. After removing the supernatant, each sample was washed once with 500 µL of FastAP buffer and incubated with 50 µL of freshly prepared alkaline phosphatase reaction mix consisting of 5 µL of 10X FastAP buffer, 2 µL of murine RNase inhibitor (40 U/µL) (NEB), 2 µL of Turbo™ DNase (Invitrogen), 3 µL of FastAP™ thermosensitive alkaline phosphatase (Thermo Fisher) (1 U/µL) and 38 µL of nuclease-free water at 37 °C for 15 min, 1200 rpm. Without removing the alkaline phosphatase reaction mix, each sample was incubated with 50 µL of freshly prepared T4 polynucleotide kinase (PNK) reaction mix consisting of 20 µL of 5X PNK buffer (pH6.5) (see Buffers, supplemental material), 2 µL of T4 polynucleotide kinase (10 U/µL) (NEB) and 28 µL of nuclease-free water at 37 °C for 20 min (interval shaking for 30 s, every 2 min). Reaction mixes were removed and each sample was washed in quick succession once with 500 µL of ice-cold wash buffer, twice with 500 µL of ice-cold high salt wash buffer and twice with 500 µL of ice-cold wash buffer.

On-bead ligation of the immunoprecipitated, cross-linked RNA was performed by adding 19 µL of freshly prepared biotinylation reaction mix consisting of 0.5 µL of 1% (v/v) Tween^®^20, 2 µL of 10X T4 RNA ligation buffer, 0.7 µL of 100% dimethyl sulfoxide (DMSO), 0.2 µL of 100 mM ATP (NEB), 0.5 µL of Cytidine-5’-phosphate-3’-(6-aminohexyl)phosphate labelled with biotin (pCp-biotin) (Jena Bioscience), 0.5 µL of murine RNase inhibitor (40 U/µL) (NEB), 8 µL of polyethylene glycol (PEG) 8000 and 6.6 µL of nuclease-free water. 1 µL of high-concentration T4 RNA ligase 1 (30 U/µL) (NEB) was added per sample and incubated at 16 °C for 2 h.

Without removing the biotinylation reaction mix, the samples were washed in quick succession three times with 500 µL of ice-cold wash buffer. The wash buffer was completely removed and each sample was resuspended in 20 µL of wash buffer. Input controls were topped to a total volume of 20 µL with wash buffer. All samples and inputs were denatured and removed from the beads by the addition of 10.5 µL of 4X NuPAGE LDS sample loading buffer (Thermo Fisher) supplemented with 285 mM DTT, 70 °C for 10 min 1200 rpm.

5 µL or 25 µL of prepared sample were loaded per well on two separate 4-15% Criterion™ TGX™ Precast Midi Protein Gels (BioRad) and run at 200 V for approximately 30 min. Transfer was performed using a TransBlot™ Turbo Transfer System (BioRad) on mixed molecular weight turbo blotting mode with Trans-Blot™ Turbo Midi 0.2 µm Nitrocellulose Transfer Packs (BioRad). From this point onwards, the two membranes were processed independently.

To detect protein signals, the low input membrane (5 µl sample) was used. Membranes were blocked in 5% milk in 1x tris-buffered saline with Tween^®^ 20 (TBST) for 30 min at RT followed by three quick TBST washes. Western Blotting was performed according to the BioRad protocol. The following primary and secondary antibodies were used for western blot analyses: PKM2 (D78A4, 1:1000 in 5% BSA in TBST), HRP-linked anti-rabbit light chain IgG (#RS3251 ECM Biosciences, 1:5000 in 5% Milk in TBST). Blots were developed with Immobilon^®^ Western Chemiluminescent HRP Substrate (Merck Millipore) or SuperSignal™ West Atto Ultimate Sensitivity Chemiluminescent Substrate (Thermo Fisher). Western blot signal detection was carried out on a ChemiDoc Go Imaging System (BioRad).

To detect RNA cross-linked to protein, the high input membrane (25 µl sample) was used. RNA was detected using the Chemiluminescent Nucleic Acid Detection Module Kit (Thermo Fisher Scientific) following manufacturer’s instructions. Detection was carried out on a ChemiDoc Go Imaging System (BioRad).

### PKM2 Wildtype and Mutant Protein Purification

Production and purification of recombinant PKM2 was performed by the EMBL Protein Production and Purification Core Facility. *Escherichia coli* BL21(DE3)-CodonPlus-RIL competent cells from Stratagene were freshly transformed with the expression plasmid (pET24a expression vectors, see supplemental material vector maps for details). ON, precultures for large scale expression were grown at 37 °C in Lennox Broth medium supplemented with 30 μg/mL kanamycin and 34 μg/mL chloramphenicol. To inoculate the large-scale expression cultures, 10 mL of preculture were added to 1 L of Terrific Broth medium supplemented with 2 mM MgSO4, 0.05% glucose, 1.5% lactose, 30 μg/mL kanamycin and 34 μg/mL chloramphenicol. Cultures were grown at 37°C until optical density (OD) at 600 nm reached around 0.6, after which the growth temperature was reduced to 18 °C. Following ON auto-expression at 18 °C, the cultures were harvested by centrifugation at 5,000 rcf for 30 min at 4 °C and the pellets were flash-frozen in liquid nitrogen and stored at -80 °C until starting the protein purification.

The cell pellet was resuspended in ice-cold protein production lysis buffer (see Buffers, supplemental material). All further purification steps were performed at 4-8 °C. Cells were lysed by five passages through a microfluidizer, followed by centrifugation at 14,000 rcf for 20 min at 4 °C. The cleared lysate was loaded onto a 5 mL Protino nickel-nitrilotriacetic (Ni-NTA) acid fast protein liquid chromatography column by Macherey-Nagel pre-equilibrated with equilibration buffer (see Buffers, supplemental material). After loading, the Ni-NTA column was washed with equilibration buffer and eluted with equilibration buffer supplemented with 250 mM imidazole. After sodium dodecyl sulfate-polyacrylamide gel electrophoresis (SDS-PAGE) analysis, elution fractions containing PKM2 were pooled, the histidine-tag was cleaved by addition of histidine-tag specific tobacco etch virus (His-TEV) protease in a ratio of 1:100 of protein to protease and dialysed overnight at 4°C against dialysis buffer (see Buffers, supplemental material). The next day, cleaved PKM2 was further purified using a combination of anion exchange and heparin chromatography. The elution fractions from the Ni-NTA purification were diluted 6-fold with 50 mM HEPES (pH 7.4) supplemented with 10% glycerol to lower the NaCl concentration to 50 mM and then loaded onto a 5 mL HiTrap™ Q HP anion exchange chromatography column by Cytiva coupled in tandem to a 5 mL HiTrap™ Heparin HP affinity column by Cytiva. After washing with ion exchange equilibration buffer (see Buffers, supplemental material), the HiTrap Q HP anion exchange chromatography column was removed and PKM2 protein was eluted from the HiTrap Heparin HP column using a gradient ranging from 50 mM NaCl to 1 M NaCl over 20 column volumes. The elution fractions containing PKM2 were pooled, concentrated to 5 mL and injected into a HiLoad™ 16/600 Superdex™ 200 pg size-exclusion chromatography (SEC) column pre-equilibrated with SEC equilibration buffer (see Buffers, supplemental material). The retrieved PKM2 elution fractions were pooled and concentrated to 6-9 mg/mL. The final PKM2 protein was aliquoted, flash-frozen in liquid nitrogen and stored at -80°C until use.

The identity of the PKM2 protein was verified by mass spectrometry and its tetrameric oligomerization state and the absence of aggregates was confirmed by size exclusion chromatography coupled with multi-angle light scattering. Appropriate stability in the storage buffer was assessed by NanoDSF analysis. For each PKM2 wildtype and mutant version, proteins were expressed and purified as independent biological triplicates.

### Electrophoretic mobility shift assay (EMSA) - non competitive

A 20 nM solution of the respective Cy5-labelled RNA (see RNA oligos, supplemental material) in 2xD buffer (see Buffers, supplemental material) was denatured at 95 °C for 2.5 min, stabilized by the addition of MgCl_2_ (final concentration of 2.5 mM) and placed on ice for 5 min.

A dilution series ranging from 0 to 30 µM of the respective recombinant PKM2 protein was prepared in PCR tubes using EMSA protein buffer (see Buffers, supplemental material; total volume: 10 µL per sample). Three independent protein preparations of PKM2 were used as biological replicates. To each sample, 1 µL of EMSA binding reaction buffer (see Buffers, supplemental material) and 10 µL of the prepared Cy5-labelled RNA were added (final RNA concentration: 10 nM; final recombinant protein concentration: 0 to 15 µM). Samples were incubated at 25 °C for 20 min in a thermocycler.

After pre-running a BioRad 4-15% Criterion™ TGX™ Precast Midi Protein gel (BioRad) for 15 - 20 min at 100 V and 4 °C in 1x native gel running buffer (see Buffers, supplemental material), 19.8 µL of the prepared samples were loaded per well and the gel was run protected from light for approximately 45 min at 100 V and 4 °C. Fluorescence signals were detected on a Typhoon™ laser scanner using 600 V and 50 µm pixel size settings on Cy5-channel. Fluorescence signals were quantified using the BioRad Image Lab software. All results shown are based on the background-corrected, adjusted volume and were normalized to the no protein control.

### Electrophoretic mobility shift assay (EMSA) - competitive

For competitive EMSAs, a fixed concentration of 7.5 µM recombinant PKM2 protein and 10 nM of Cy5-labelled RNA was used. During the binding reaction, 0-50 µM of Pyruvate (Sodium pyruvate, P2256, Sigma-Aldrich), 0-50 µM Phosphoenolpyruvate (Phospho(enol)pyruvic acid monopotassium salt, P7127, Sigma-Aldrich) or 0-25 µM FBP (D-Fructose 1,6-bisphosphate trisodium salt hydrate, F6803, Sigma-Aldrich) were supplemented. For details on the protocol see *Electrophoretic mobility shift assay (EMSA) - non competitive*.

### Nano differential scanning fluorimetry

All buffers were double-filtered and degassed prior to use. The recombinant PKM2 variants were diluted to a final concentration of 3 μM in 19 μL of NanoDSF/MP protein buffer (see Buffers, supplemental material) or PBS substituted with 2.5 mM MgCl2. 1 μL of either RNase-free water or FBP (F6803, Sigma-Aldrich) was added to a final concentration of 0.5 mM. Samples were incubated at 25 °C for 20 min. Samples were loaded in Standard nanoDSF grade capillaries (NanoTemper Technologies) and measured on a Prometheus NT.48 with backscattering optics (NanoTemper Technologies) applying 80% excitation power and a temperature gradient ranging from 25 °C to 95 °C, increasing by 1 °C per min. Melting temperatures ( i.e. inflection points for F330nm) were calculated with the PR. ThermControl software (NanoTemper Technologies). Each experiment was conducted in technical duplicates, with three biological replicates of each condition using three independently produced and purified recombinant human PKM2 preparations per variant.

### Mass photometry

All buffers were double-filtered and degassed prior to use. The recombinant PKM2 variants were pre-diluted to a final concentration of 3.2 μM in NanoDSF/MP protein buffer (see Buffers, supplemental material) or PBS substituted with 2.5 mM MgCl2. 1.25 μL of each pre-dilution were combined with 1.25 μL of the respective buffer and either 2.5 μL of 1 mM FBP (F6803, Sigma-Aldrich) diluted in the respective buffer or 2.5 μL of the respective buffer (final PKM2 concentration 800 nM, final FBP concentration 500 μM). The samples were incubated at 25 °C for 20 min. Immediately before measuring each sample, was first diluted 1:1 in the respective buffer and then loaded on the instrument as a 1:20 dilution in the respective buffer (final PKM2 concentration 20 nM, final FBP concentration 12.5 μM). Each experiment was conducted in single measurements using a Two^MP^ mass photometer (Refeyn Ltd.). Videos of 1 min were recorded with normal mode and regular image size using the Acquire^MP^ version 2024 R2.1 software (Refeyn Ltd). The data were analyzed using the Discover^MP^ version 2024 R2.1 software (Refeyn Ltd). Bovine serum albumin (BSA) and immunoglobulin G (IgG) were used to generate the standard contrast-to-mass calibration curve. Biological triplicates were generated using three independently produced and purified recombinant protein preparations.

### Pyruvate kinase activity assay

Pyruvate kinase activity levels of PKM2 variants were assessed using a pyruvate kinase assay kit (Ab83432, Abcam) following manufacturer’s instructions. In order to stay within the kit’s linear capacities, 5 ng of wild-type PKM2, 500 ng of MutA and 1 ng of PXX3, K433A, R436A and R455A were used, respectively. Each sample was supplemented with FBP (F6803, Sigma-Aldrich) (final concentration: 0 μM, 0.1 μM or 500 μM) in kit assay buffer (total volume: 50 µl). The reaction was started by providing each sample with 50 μL of pyruvate kinase reaction mix consisting of 44 μL of kit assay buffer, 2 μL of substrate mix, 2 μL of enzyme mix and 2 μL of OxiRed^TM^ probe. The optical density (OD) was measured at 570 nm every min for 40 min on a Tecan Spark® plate reader. For each biological replicate, pyruvate kinase activity was determined using the formula provided by the manufacturer’s protocol:

*PK activity* = (*B* x *Sample Dilution Factor*)/((*T*2−*T*1) x *V*)

*B:* amount of pyruvate derived from linear regression of pyruvate standard curve and insertion of the measured difference in OD between timepoints T2 and T1 into the regression equation [nmol], *T:* Chosen timepoints within linear reaction range [min], *V:* sample volume [mL].

Each experiment was conducted in biological triplicates deriving from three independently produced and purified recombinant protein preparations per sample condition, respectively. Each biological replicate was measured in technical duplicates.

### Quantification and Statistical Analysis

Statistical analysis was performed by unpaired or paired Student’s t test or two- or one-way ANOVA without correction for multiple comparison (Fisher LSD test). Please see figure legends for detailed information. Significance levels were set at p* < 0.05, p** < 0.01 and p*** < 0.001. For statistical analysis, GraphPad Prism was used.

## ACKNOWLEDGEMENTS

We thank current and former members of the Hentze laboratory for their suggestions. We thank S. Colucci for advice on the EMSA experiments. We acknowledge the EMBL core facilities for Protein Expression and Purification, and Metabolomics for their expert services, with special thanks to K. Remans and J. Scheurich, and the Centre of Biophysics of Macromolecules and their Interactions (PFBMI), Institut Pasteur, Paris. This project has received funding from the Peter und Traudl Engelhorn Stiftung, the European Commission (under the Marie Skłodowska-Curie grant agreement no 101102982) and the Christiane Nüsslein-Volhard Foundation (P.S.). M.W.H. gratefully acknowledges funding from the EMBL Technology Development Fund as well as support by the Manfred-Lautenschläger Foundation. This project has received funding from the European Union’s Horizon 2020 research and innovation program under grant agreement No 101004806.

## AUTHOR CONTRIBUTIONS

Conceptualization, P.S. and M.W.H.; Methodology, P.S., C.S., K.L., A.B., and D.F.-A.; Investigation, P.S., C.S., and K.L.; Writing - Original Draft, P.S. and M.W.H.; Writing - Review and Editing, P.S., C.S., K.L., A.B., D.F.-A, and M.W.H; Supervision, P.S. and M.W.H.

## DECLARATION OF INTERESTS

The authors declare no competing interests.

## SUPPLEMENTAL ITEM TITLES

SUPPLEMENTAL MATERIAL: Buffers, RNA oligos, Vector Maps

## EXTENDED DATA FIGURES

**Extended Data Fig 1:**
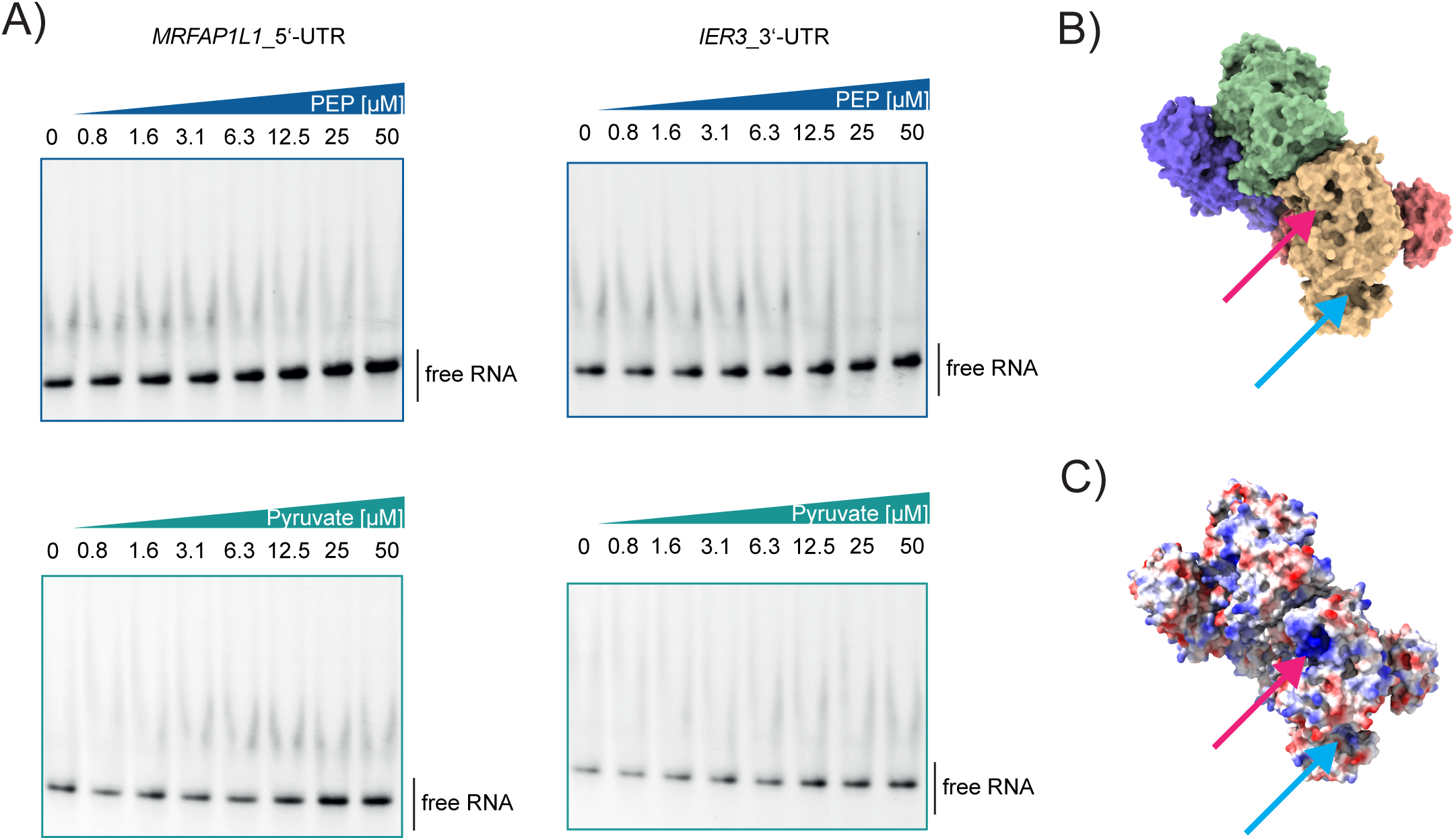
A positively charged FBP-associated surface mediates PKM2 RNA binding. (A) EMSA analysis of recombinant WT PKM2 and Cy5-labelled PKM2 RNA target oligos *MRFAP1L1*_5’-UTR (45mer) and *IER3*_3’-UTR (40mer) ^30^ in competition with its substrate phosphoenolpyruvate (PEP) and its product pyruvate. N=3 (for each biological replicate an independent recombinant protein preparation was used). Representative EMSAs are shown. (B) Structure of tetrameric human WT PKM2 in complex with FBP. Each PKM2 subunit is shown in surface mode in differing colors. The red arrow indicates the positive patch encompassing positive amino acid residues K433, R436 and R455; the blue arrow indicates the catalytic site, in the yellow PKM2 subunit, respectively. Image was created using ChimeraX and the x-ray diffraction derived PKM2 crystal structure 1T5A from the RCSB Protein Data Bank (https://doi.org/10.2210/pdb1t5a/pdb) ^12^. (C) Identical to Extended Data Fig 1B, color surface by electrostatic potential.

**Extended Data Fig 2:**
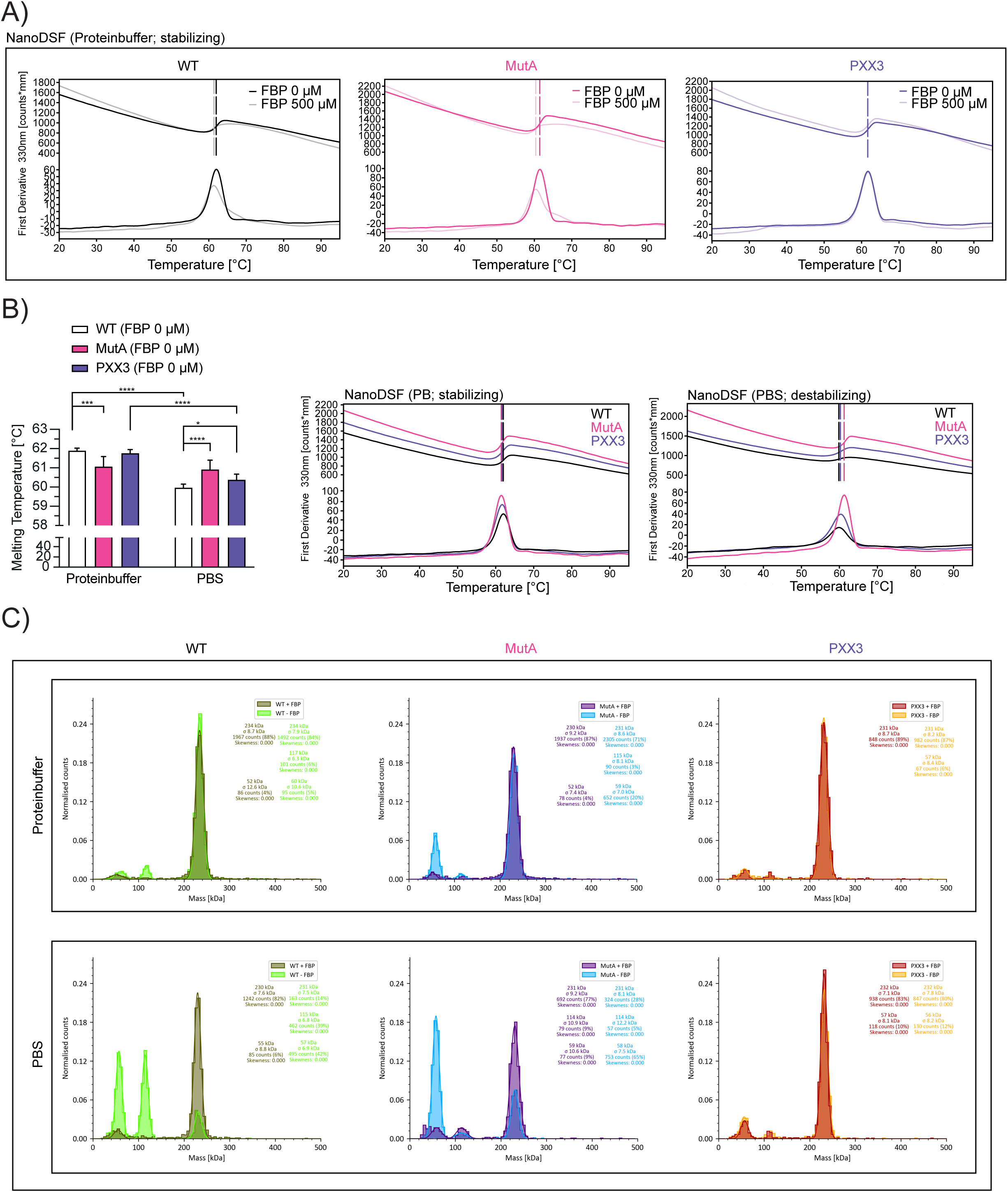
PKM2 RNA binding requires an intact FBP-responsive surface and is favored by the tetrameric state. (A) Representative NanoDSF thermograms of 3 µM PKM2 WT, MutA and PXX3 ±500 µM FBP in protein buffer. N=3 (for each biological replicate an independent recombinant protein preparation was used). (B) Quantification of melting temperatures of PKM2 WT, MutA and PXX3 ±500 µM FBP in protein buffer or PBS +2.5 mM MgCl_2_. N=3 (for each biological replicate an independent recombinant protein preparation was used). Mean +SD. One-way ANOVA. *p < 0.05; **p < 0.01; ***p < 0.001; ****p < 0.0001. Overlay of NanoDSF thermograms of PKM2 WT, MutA and PXX3 are shown (see also Extended Data Fig 2A) (C) Representative mass photometry histograms of PKM2 WT, MutA and PXX3 in protein buffer or PBS +2.5 mM MgCl_2_ ±FBP.

**Extended Data Fig 3:**
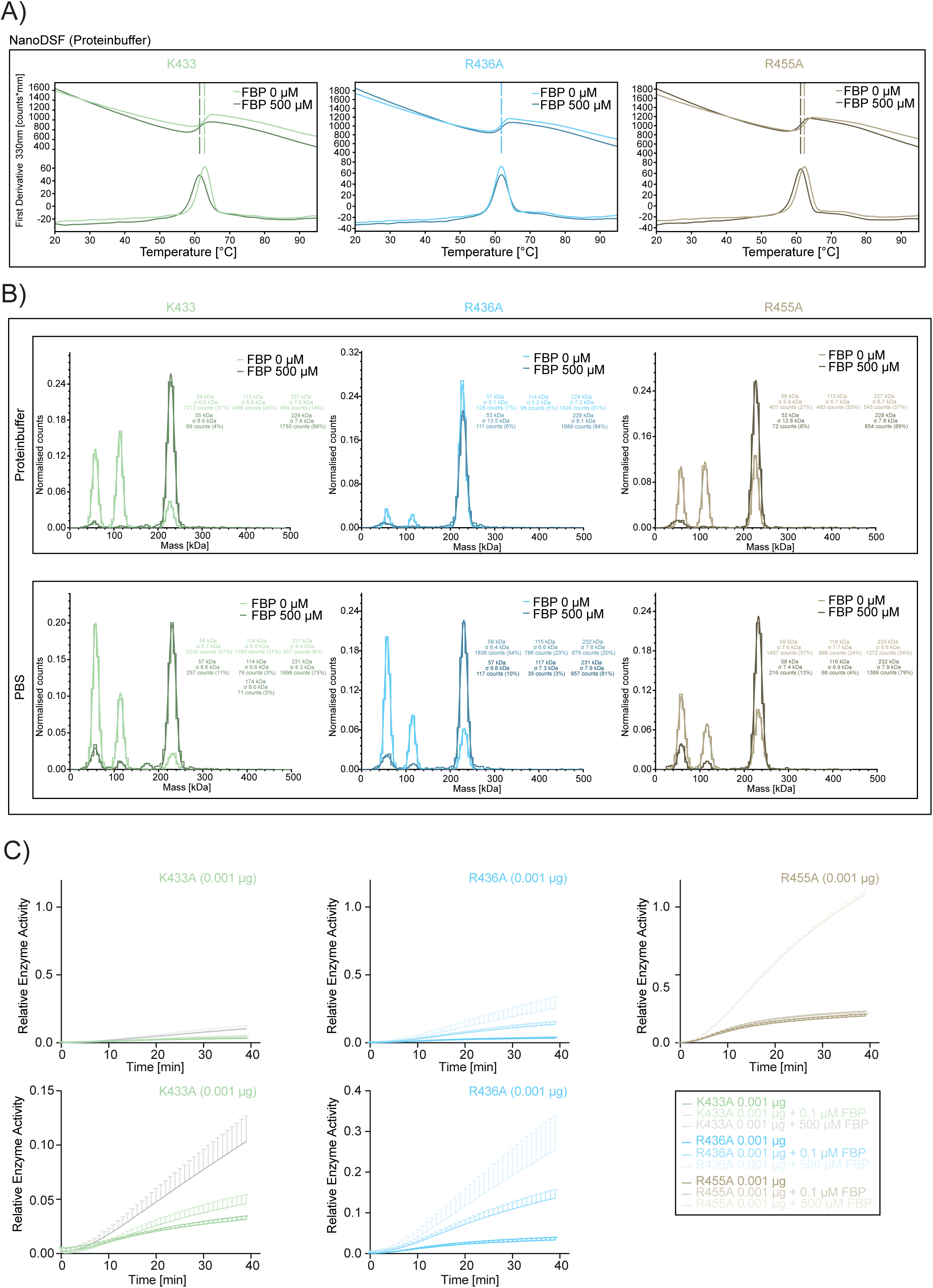
Single-residue mutations uncouple PKM2 RNA binding, FBP responsiveness and enzymatic activity. (A) Representative NanoDSF thermograms of 3 µM K433A, R436A and R455A ±500 µM FBP in protein buffer. N=3 (for each biological replicate an independent recombinant protein preparation was used). (B) Representative mass photometry histograms of K433A, R436A and R455A in protein buffer or PBS +2.5 mM MgCl_2_ ±FBP. (C) Pyruvate kinase activity curves ±FBP. In order to stay within the assay kit’s linearity capacities, 0.001 µg K433A, R436A and R455A were used, respectively. Different amounts of recombinant protein were taken into account and corrected during calculation of relative enzyme activity. N=3 (for each biological replicate an independent recombinant protein preparation was used). Mean +SD.

## Notes

### Competing Interest Statement

The authors have declared no competing interest.

